# The trade-off between pulse duration and power in optical excitation of midbrain dopamine neurons approximates Bloch’s law

**DOI:** 10.1101/2021.04.22.440938

**Authors:** Vasilios Pallikaras, Francis Carter, David Natanael Velázquez Martínez, Andreas Arvanitogiannis, Peter Shizgal

## Abstract

**Background:** Optogenetic experiments reveal functional roles of specific neurons. However, such inferences have been restricted by widespread adoption of a fixed set of stimulation parameters. Broader exploration of the parameter space can deepen insight into the mapping between selective neural activity and behavior. In this way, characteristics of the activated neurons, such as temporal integration, can be inferred.

**Objective:** To determine whether an equal-energy principle accounts for the interaction of pulse duration and optical power in optogenetic excitation.

**Methods:** Six male TH::Cre rats worked for optogenetic (ChannelRhodopsin-2) stimulation of Ventral Tegmental Area dopamine neurons. We used a within-subject design to describe the trade-off between pulse duration and optical power in determining reward seeking. Parameters were customized for each subject on the basis of behavioral effectiveness.

**Results:** Within a useful range of powers (~12.6-31.6 mW) the product of optical power and pulse duration required to produce a given level of reward seeking was roughly constant. Such reciprocity is consistent with Bloch’s law, which posits an equal-energy principle of temporal summation over short durations in human vision. The trade-off between pulse duration and power broke down at higher powers.

**Conclusions:** Optical power can be substituted for pulse duration to scale the region of neuronal excitation in behavioral optogenetic experiments. Power and duration can be adjusted reciprocally for brief durations and lower powers. The findings demonstrate the utility of within-subject and trade-off designs in optogenetics and of parameter adjustment based on functional endpoints instead of physical properties of the stimulation.

**Highlights:**

- We provide behaviorally derived intensity-duration curves for ChannelRhodopsin-2.
- Duration trades off almost perfectly with power within useful ranges.
- This trade-off breaks down at high optical powers.
- Pulse duration and optical power scale the area of neuronal excitation equivalently.
- Behaviorally derived trade-offs can reveal optogenetic excitation mechanisms.

## Introduction

Optogenetic excitation establishes causal links between activation of specific neurons and behavior^1,2^. However, researchers applying optogenetic manipulations face the practical problem of parameter selection. Using parameters already identified in published studies provides a sub-optimal solution to this problem. Indeed, the widespread adoption of a fixed set of stimulation parameters by the optogenetics community leaves much of the stimulation-parameter space unexplored, the mapping between parameter values and observed behavior partly unexplained, and individual differences in opsin expression and probe placement unaddressed. To realize the full potential of optogenetics, we must map observable behavior onto stimulation parameters for each subject. We do so here for optical self-stimulation (oICSS)^3–6^. Rats were trained to work for optical stimulation of Channelrhodopsin-2 (ChR2)-expressing dopamine neurons in the Ventral Tegmental Area (VTA). We determined how pulse duration and optical power interact to control reward seeking, and we assessed the correspondence between this interaction and Bloch’s law^7,8^, a principle of temporal summation in visual perception.

The irradiance produced by an optical pulse of a given duration, and hence the amplitude of the induced photocurrent, decays as a function of distance from the tip of the implant^9^. Consequently, the greater the distance between an opsin-expressing neuron and the tip of an optical fiber, the longer the required duration of a pulse to trigger an action potential^10^. Thus, at a given optical power, the cross-sectional area of the region wherein a pulse activates opsin-expressing neurons scales with pulse duration. In this sense, pulse duration and optical power codetermine the area of excitation and the number of recruited neurons.

Dopamine release in the Nucleus Accumbens (NAc) driven by optical stimulation of midbrain dopamine neurons increases as a function of optical power^11^. Operant performance for such stimulation increases as a function of both optical power^6,12^ and pulse duration^12^. However, the form of the interaction between optical power and pulse duration in determining the behavioral effectiveness of the stimulation has not been described empirically. Such strength-duration curves have been derived theoretically for ChR2-expressing neurons^10,13^, but, to our knowledge, no such curves are available either for activation of midbrain dopamine neurons or for oICSS of these neurons. We provide such curves here for the case of oICSS.

In human vision, there is a simple relationship between the intensity and duration of a light flash required to produce a just-detectable stimulus. As flash duration increases, the required illuminance declines initially as a power function of duration and is then roughly constant beyond a critical duration that depends on the spatial frequency^14^. This relationship is called Bloch’s law^7,8^. Here, in the case of ChR2-mediated activation of midbrain dopamine neurons, we determine whether pairs of pulse durations and optical powers that cause equivalent reward seeking approximate Bloch’s law. This question is germane to the development and use of optogenetic prostheses and to linking specific neural populations to function.

**Table 1.**
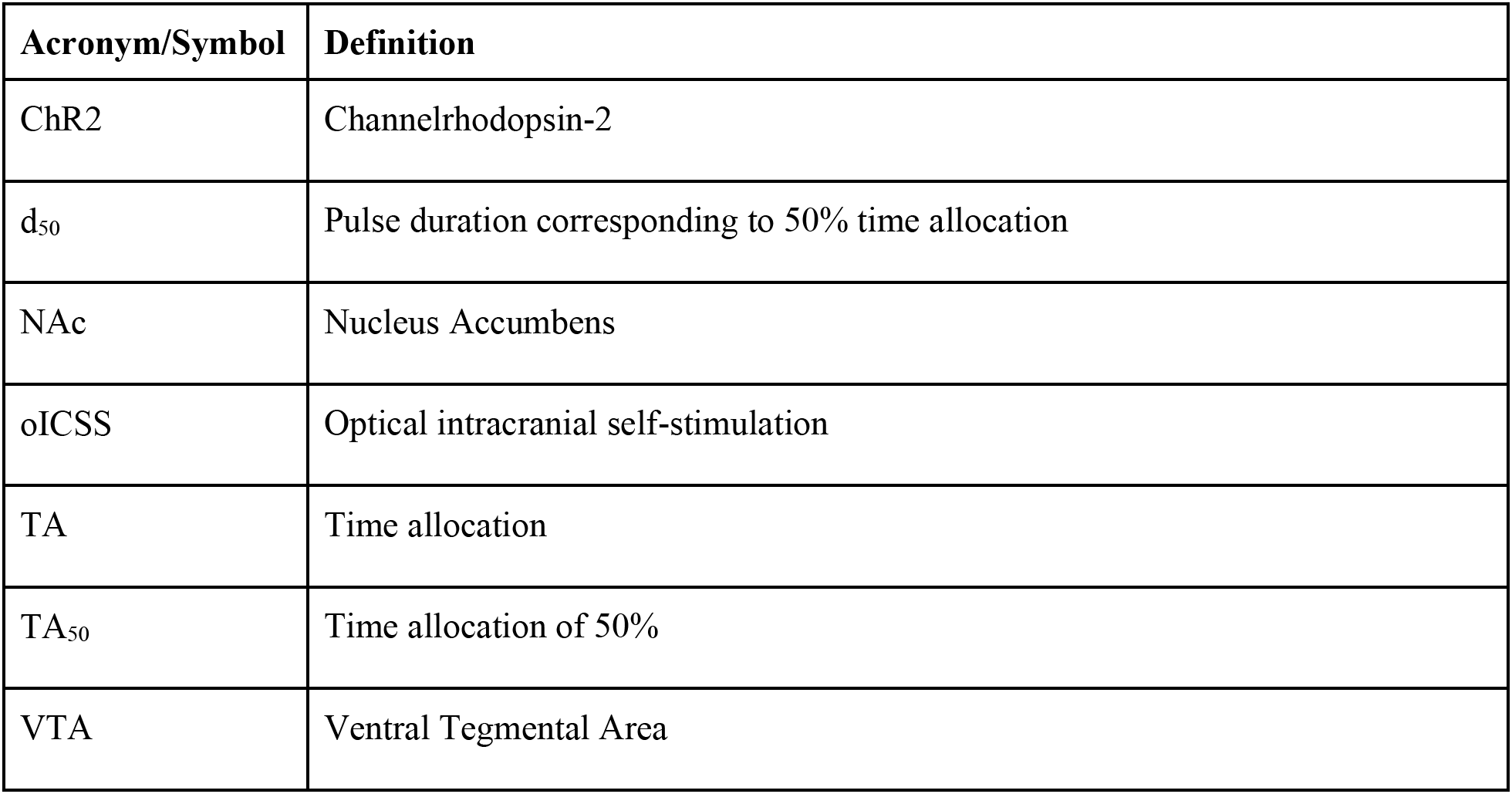
Definitions of acronyms and symbols used.

## Methods

### Subjects

Subjects were 6 male, TH::Cre, Long-Evans rats weighing ~350-475 g at surgery. Rats were singly housed on a reverse 12 h-light cycle and had free access to chow (Envigo #2014). Subjects deemed overweight for their age^15^ were put on a weight-maintenance feeding schedule. Subject ELOP18 had bilateral implants and contributed individual datasets for each hemisphere. Procedures for animal care were approved by Concordia University Animal Research Ethics Committee (Protocol #: 30000302) and adhered to standards of the Canadian Council on Animal Care.

### Surgery

Surgical procedures are described in detail elsewhere^6^. Rats were anesthetized (IP ketamine; 87 mg/kg/xylazine; 13 mg/kg) and positioned in a stereotaxic frame for injection of the viral vector (AAV5-DIO-ChR2-EYFP; University of North Carolina) via a microinfusion pump (Kent Scientific). The injector was aimed at the VTA (AP: −5.5 & −6.2, ML: ± 0.7, DV: −7.4 & −8.4), and 1 μl of virus was infused at each site at a rate of 0.1 μl/min. Optical fiber implants (300 μm diameter, 0.39 NA) were chronically positioned above the VTA (AP: −5.8, ML: ± 0.7, DV: −7.7).

### Apparatus

We used six operant boxes equipped with a house light and a retractable lever with a cue light above it. The house light flashed once per second during the inter-trial interval (ITI), and the cue light was illuminated while the lever was depressed. The laser head (462 nm; SLOC) was connected via a fiber-optic cable (300 μm diameter, 0.39 NA) to a single-channel rotary joint (FRJ_1×1_FC-FC; Doric Lenses) mounted above the box. An optical fiber cable^16^ (300 μm diameter, 0.39 NA) was connected to the implant via a ceramic sleeve. We set optical power with a power meter (PM100D; Thorlabs) by measuring the continuous-wave output of the lasers prior to each session.

### Self-Stimulation Training

Four weeks post-surgery, we used successive approximation to train subjects to perform oICSS under an FR-1 schedule. Stimulation trains, 1 s in duration, consisted of 5 ms pulses delivered at 40 Hz. Optical power ranged from 20-40 mW.

### Cumulative hold-down schedule of reinforcement

Subjects were trained to hold down an operant lever to trigger stimulation^17^. A pulse train was delivered each time the cumulative hold-down time reached 2 s. After stimulation was triggered, the lever was retracted for 1.5 s, and the trial timer was paused until the lever re-extended. The dependent variable was time allocation: the proportion of total trial time spent working. Pauses between lever presses shorter than 1 s were classified as work. During such instances, the rat typically holds its paw on or above the lever^17^.

### Pulse Duration Sweeps

A sweep was a set of 10 trials over which the pulse duration was decreased systematically (“swept”) while all other parameters were held constant. Trials lasted for 50 s and were preceded by a 10 s ITI. At the 8th second of the ITI, a train of priming stimulation was delivered. This stimulation was identical to the train available on the first trial of the sweep. The pulse duration was fixed within trials. On the first two trials of each sweep, pulse duration was set to the longest value tested and then reduced in eight proportional steps from the 3rd to the 10th trial. The range of pulse durations was customized for each rat and condition. Sessions, roughly 2 h in duration, consisted of 10 pulse-duration sweeps. The first sweep and the first trial of every sweep were considered warm-ups, and their data were discarded. We tested conditions at various optical powers (between 10 mW - 50 mW) at each stimulation site. Five sessions (45 sweeps) were run in every optical power condition. All data was analyzed in Matlab (MathWorks).

### Curve Fitting

Due to noise in the lateral position of curves, plots of averaged time allocation versus pulse duration have a shallower slope than the curves obtained on individual sweeps^18^. Therefore, we fit a function to the time allocation values for each sweep and averaged its parameters^18^.

Although time allocation varied systematically with pulse duration on the great majority of sweeps, some aberrant cases were observed. We filtered the data by eliminating sweeps in which the average time allocation on the first four trials did not exceed the average on the last four trials by at least 20%. Sensitivity to outliers was reduced by a robust fitting method based on Tukey’s bisquare estimator^19^.

The following four-parameter equation was fitted:

(*TA - TA_min_*) / (*TA_max_ - TA_min_*) = 1 / [1 + exp{-*slp* × (log_10_ (*d*) - *loc*)}]

Where:

*d*: Pulse duration.

*loc*: Value of the location parameter.

*TA*: Time allocation.

*TA_min_*: Minimal time allocation.

*TA_max_*: Maximal time allocation.

*slp*: Slope parameter determining the steepness of the rise.

### Histology

Rats were anesthetized (sodium pentobarbital; 200 mg/kg IP) and perfused transcardially using saline and 4% paraformaldehyde. Brains were post-fixed for 24 hours in 4% paraformaldehyde and subsequently cryoprotected by successive immersion in phosphate-buffered solutions of 15% and 30% sucrose until they sank. Brains were then stored at −80° C. Using a cryostat, 40 μm sections were cut through the VTA and mounted on glass slides. Slices were stained with DAPI (ThermoFisher). Native fluorescence of eYFP revealed expression of ChR2 in dopamine neurons. Figure 1 shows implant placements.

**Figure 1.**
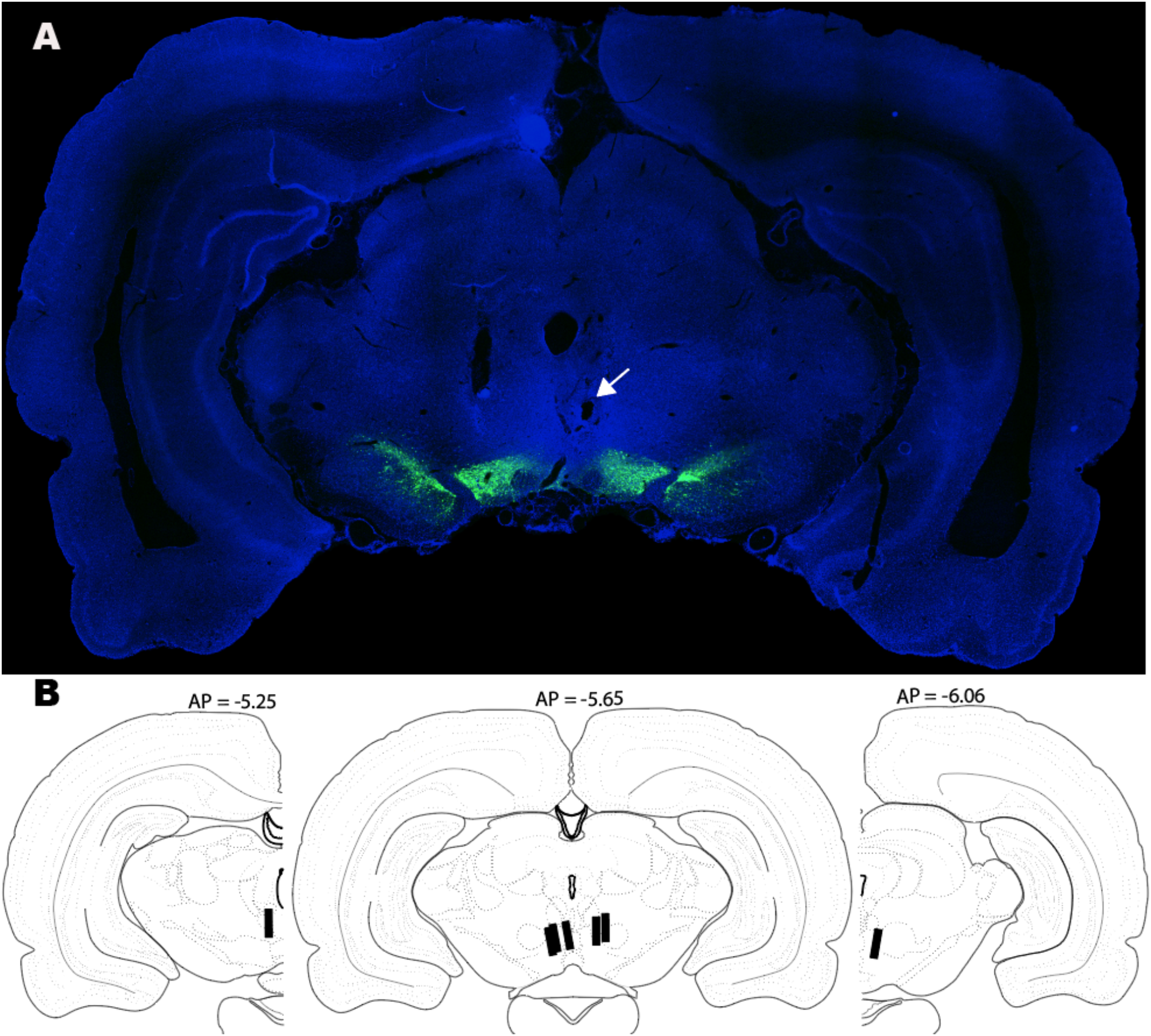
**A:** Histological image for subject OP13; eYFP expression (green) shown along with DAPI (blue) for anatomical reference (image enhanced by rescaling the distribution of pixel intensities). Estimated fiber-tip location: white arrow. **B:** Black lines indicate implant placements. Adapted from Swanson^20^.

## Results

### Time Allocation as a Function of Pulse Duration

Time allocation grew as a sigmoidal function of pulse duration in all subjects. Panel A of Figure 2 depicts single-session data from an exemplar subject (OP13). The average time-allocation values for this condition are shown in panel B (green diamonds). Also shown in panel B is a curve obtained by averaging the parameters of sigmoidal functions fitted to the data from the individual sweeps for this condition^18 (Fig. 14)^. The shaded band reflects the 95% confidence interval around the location parameter. ***TA_50_***, refers to time allocation of 50%, and ***d_50_*** refers to the corresponding pulse duration. The impact of the curve-fitting procedure on noisier data is shown in appendix A (Figure A.1). Appendix B (Tables B.1-5) provides fitted parameter values for the dataset.

**Figure 2.**
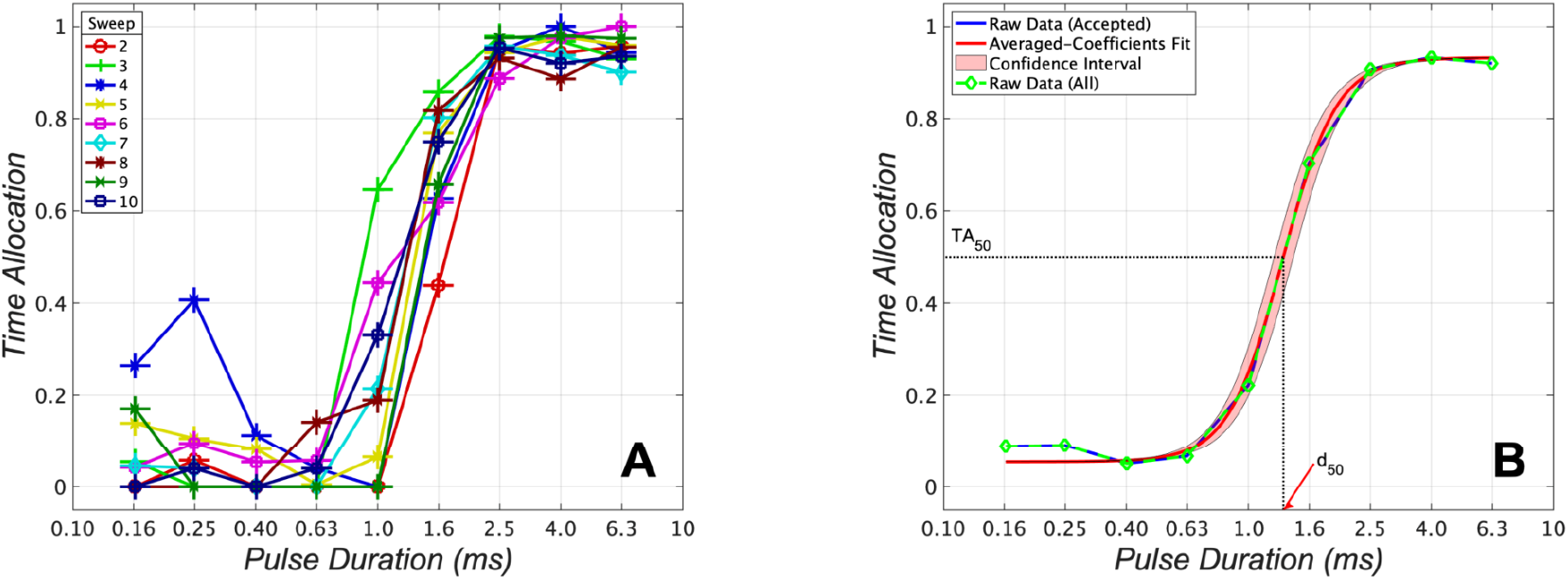
Pulse-duration sweep data for subject OP13 (optical power: 31.6 mW). **A:** Single-session data depicting individual time-allocation-versus-pulse-duration curves. **B:** Conventional averaging of time allocation over five test sessions (45 sweeps; green line) and curve produced by averaging the parameters of fitted sigmoidal functions (red line) with 95% confidence interval surrounding the location parameter (pink).

### Pulse Duration Trades off with Optical Power

We determined how optical power trades off against pulse duration to hold time allocation constant. Increasing the optical power systematically shifted the time-allocation curves leftwards along the pulse-duration axis (Figure 3). Thus, the same level of time allocation can be achieved by a stimulation train composed of brief, high-power pulses and a train composed of longer, lower-power pulses. However, at 6/7 stimulation sites that could be tested successfully at the highest optical power (50 mW at 4 sites; 44.6 mW at 1 site; 31.6 mW at 1 site), the final increment failed to produce a leftward shift. Thus, there was an upper limit (generally, beyond 31.6mW) on the optical power that trades off against pulse duration to hold behavior constant.

**Figure 3.**
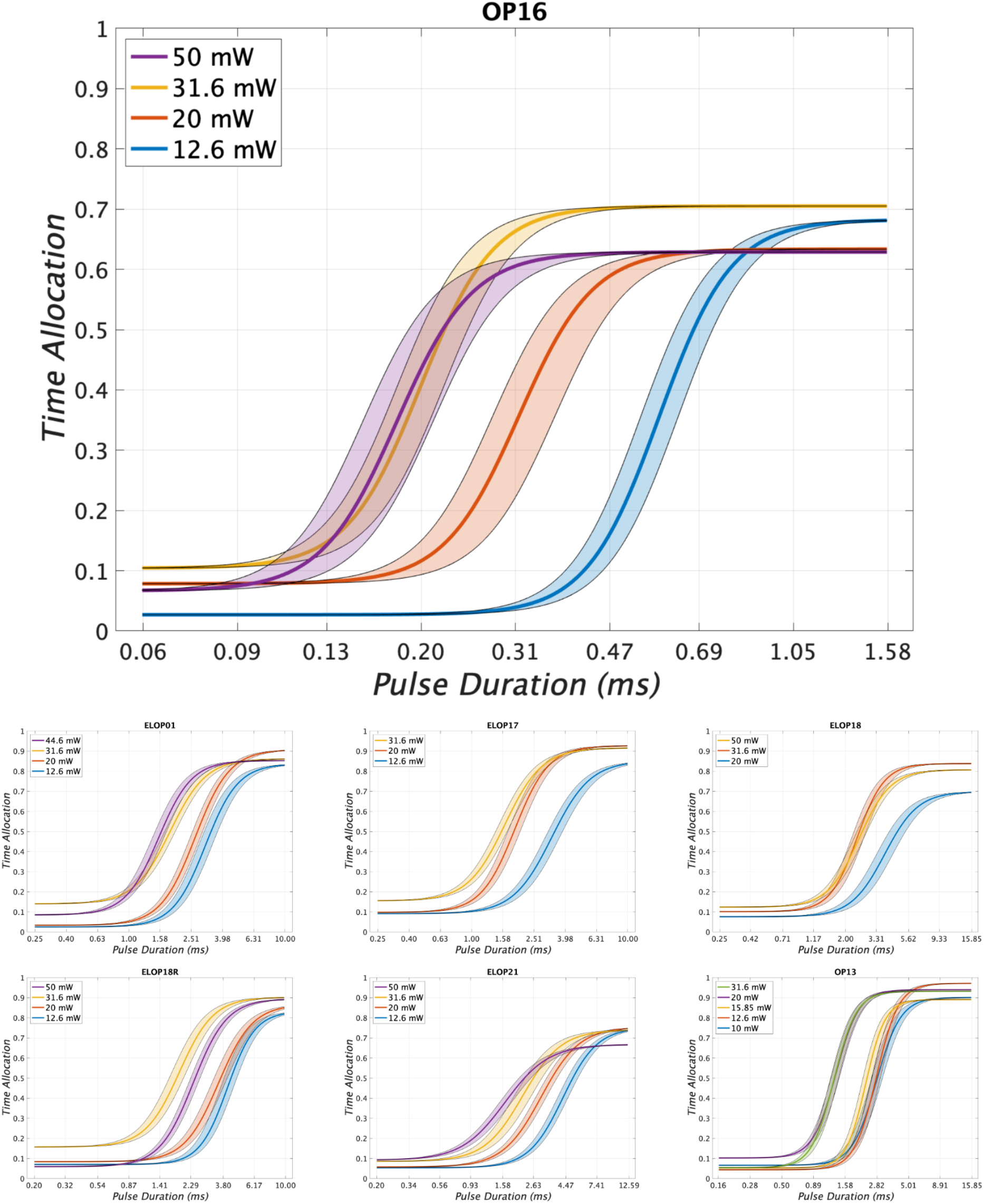
Fitted time-allocation-versus-pulse-duration curves for each optical power at all 7 stimulation sites. Over a range of powers, there is a systematic trade-off between pulse duration and optical power in maintaining half-maximal time allocation. At 6 of 7 sites, the observed trade-off breaks down at the highest power increment (31.6-50 mW at 4 sites; 31.6-44.6 mW at 1 site; 20-31.6 mW at 1 site).

### The Trade-off Between Pulse Duration and Optical Power Approximates Bloch’s Law

We determined whether the trade-off between the optical power and pulse duration (***d_50_***) required to hold time allocation constant at ***TA_50_*** corresponds to Bloch’s Law. If Bloch’s law holds, constant optical-energy deposition will produce a constant level of behavior (time allocation) over an initial range of pulse durations. Thus, the product of optical power and ***d_50_*** will be constant over an initial range of pulse durations.

To assess the correspondence of the data to Bloch’s law, we mean-centered the common logarithms of the ***d_50_*** values and plotted them against the common logarithm of the optical power (Figure 4A) and the product of power and ***d_50_*** (Figure 4B). The data from the highest optical powers tested were excluded. Perfect reciprocity between pulse duration and optical power holds if the slope of these lines is −1 (Figure 4A) and 0 (Figure 4B), respectively. The 95% confidence intervals surrounding the slopes of the lines of best fit include −1 and 0, respectively; −1.017 ± 0.174 (Figure 4A), 0.005 ± 0.048 (Figure 4B). Figure A.2 shows an analogous plot with no data excluded.

**Figure 4.**
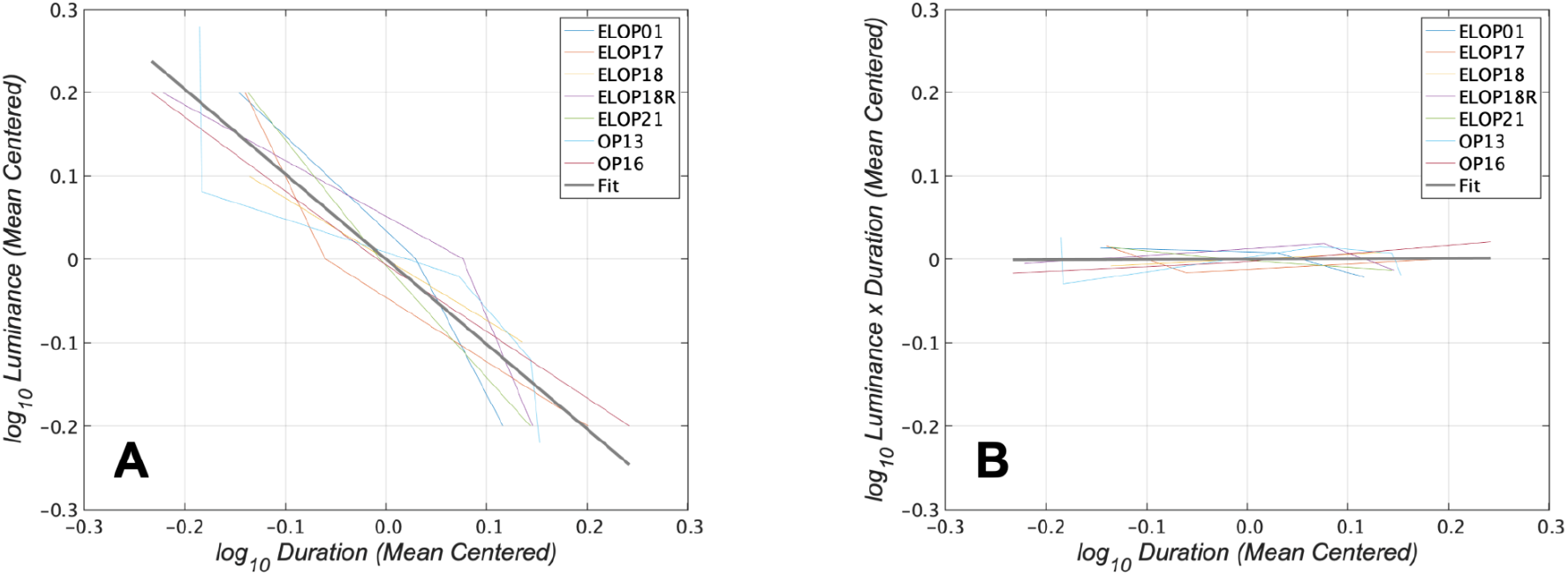
The trade-off between pulse duration and optical power approximates Bloch’s law. The data from the highest optical powers tested were excluded. **A:** Mean-centered optical-power-vs-pulse-duration curves for individual stimulation sites (colored lines) and regression line (thicker black line; slope; −1.017 ± 0.174. **B:** Mean-centered energy (optical power × pulse duration) versus pulse-duration curves for individual stimulation sites (colored lines) and regression line fitted to the entire dataset (thicker black line; slope 0.005 ± 0.048).

## Discussion

Only a restricted region of the parameter space for optogenetic stimulation has been explored comprehensively in behavioral optogenetic studies. To better understand how stimulation parameters interact in the activation of opsin-expressing neurons and translate into behavior, we measured the trade-off between optical power and pulse duration in optical self-stimulation of ChR2-expressing midbrain dopamine neurons.

To our knowledge, this is the first behavioral experiment to describe intensity-duration curves for ChR2. The intensity-duration trade-off documented here implies that the pulse duration scales the number of stimulated opsin-expressing neurons and thus provides a convenient means of achieving such scaling in optogenetic setups requiring manual control of optical power. The findings highlight the usefulness of within-subject and behavioral trade-off designs in optogenetic experiments and the utility of testing parameter ranges well beyond the values that have become an informal standard in behavioral optogenetic studies.

### How Optical Power and Pulse Duration Codetermine the Number of Stimulated Dopamine Neurons

The key causal event in this study is the firing of ChR2-expressing dopamine neurons. The following reviews how this event was inferred from behavior.

For simplicity, we assume initially equal expression of ChR2 in dopamine neurons and equal excitability across neurons and pulses. Under these conditions, there is a sharp boundary dividing the region where a pulse is too weak to trigger action potentials and the region where a pulse always triggers a spike. Relaxing these assumptions creates multiple boundaries (e.g., one for each subpopulation defined on the basis of excitability) and renders these boundaries fuzzy in time and space, but it does not change the fundamental point: pulse duration acts as a spatial variable.

Irradiance is highest at the fiber tip^9^. Thus, the induced photocurrent rises most steeply and surpasses the threshold for triggering an action potential soonest in the nearest opsin-expressing neurons. As distance from the tip increases, irradiance and the rise in the resulting photocurrent fall off, delaying spike onset. At the sharp boundary defined by the simplified assumptions, the spike is initiated only at the very end of the pulse. Increasing the pulse duration pushes the boundary outwards by providing more time for the photocurrent to achieve suprathreshold depolarization. Thus, pulse duration trades off against the other parameter that acts spatially: the optical power. Bloch’s law asserts that this trade-off is reciprocal over an initial range of durations (i.e., the product of the two variables is constant over this range). Figure 4 shows such reciprocity.

The pulse duration was swept from trial to trial within each test session. Within the swept range is ***d_50_***: the pulse duration required to produce 50% time allocation. Figure 3 shows that changing the optical power shifts ***d_50_***. We next show that given the above assumptions and a long-standing account of how stimulation-induced firing is linked to ICSS performance, the boundary of the region containing the optically excited dopamine neurons is the same for each pair of optical powers and ***d_50_*** values. If so, the same number of dopamine neurons are excited. Given that the pulse frequency and train duration were constant, the aggregate rate of induced firing is constant as well.

### From Excitation of Dopamine Neurons to Behavior and Back

According to the “counter model” of spatiotemporal integration in brain reward circuitry^21–23^, the intensity of brain stimulation reward is a function of the aggregate firing rate induced by a pulse train of fixed duration. This model has been tested most extensively in the case of electrical ICSS^23^, but it is also consistent with oICSS data^6,12^. The counter model implies that beyond the outputs of the activated midbrain dopamine neurons, there is a neural signal, (dubbed “reward intensity”) whose amplitude in response to a pulse train of a given duration is a monotonic function of the aggregate firing rate^23^. According to the single-operant matching law^6,24–26^, time allocation is a monotonic function of reward intensity and hence of the aggregate firing rate as well. Thus, during a pulse-duration sweep, time allocation is a monotonic function of pulse duration. If so, every pair of optical powers and ***d_50_*** values generates the same aggregate firing rate by activating the same dopamine neurons, and the behaviorally derived trade-off function applies to the excitation of these dopamine neurons. This logic is analogous to that underlying the remarkable correspondence between the absorption spectrum of human rhodopsin and the appropriately corrected sensitivity function for scotopic vision^27^.

### Correspondence to Bloch’s Law

Both the Bunsen-Roscoe law of photochemistry^28,29^ and Bloch’s law^7,8,30^ reflect a constant-energy principle: for sufficiently short durations, the effectiveness of a light pulse is determined by the product of optical power and pulse duration. Within a range of optical powers (12.6 mW - 31.6 mW) and pulse durations (0.22-5.00 ms), our results approximate the predicted reciprocity. This implies that at least within the aforementioned limits, the constant-energy principle applies to the optical excitation of ChR2-expressing midbrain dopamine neurons.

The correspondence between the data and Bloch’s law broke down or was moderated at the highest optical powers tested. The sigmoidal curve either failed to shift appreciably (OP13, OP16, ELOP01, ELOP18), shifted too little in the leftward direction (ELOP21), or shifted rightwards (ELOP18R). The final increment in optical power should have pushed the excitation boundary outwards by the same amount as the other increments (which were of the same magnitude). The failure of the curves to shift sufficiently, at all, or in the predicted direction, implies that neurons were subtracted from the activated population in sufficient numbers to decrease, cancel, or reverse the contribution of the firings added at the periphery of the field.

How might the final increment in optical power have subtracted firings? A heating-induced decrease in neural excitability is an unlikely explanation because heating obeys an equal-energy principle analogous to Bloch’s law^31^. Moreover, the range of ***d_50_*** values was roughly an order of magnitude lower in subject OP16 than in the other subjects. The heat deposited by the parameters corresponding to ***TA_50_*** in this rat should have been an order of magnitude lower than in the other subjects, but the equal-energy principle broke down in this rat at the highest optical power tested, as it did in the others.

The highest optical power tested in this study was 50 mW, which generates an optical-power density at the fiber tip of 707 mW/mm^2^. This value greatly exceeds those used typically in determining the photocycle of ChR2. For example, the optical-power density of pulses delivered in the ChR2 photocycle study by Kuhne and colleagues^32^ was 1 mW/mm^2^, and 1-5 mW/mm^2^ is typically taken as the threshold for opening ChR2 channels^1^. We wonder whether the greatly suprathreshold optical-power densities might somehow reduce firing in the region closest to the fiber tip, thereby offsetting firings added at the periphery of the field and violating reciprocity. Is such a reduction seen in response to greatly suprathreshold irradiance in ChR2-expressing dopamine neurons observed by electrophysiological means? Such direct electrophysiological measurements would provide a strong test of the inferences drawn here concerning the application of the constant-energy principle to optogenetic stimulation.

### Implications

The demonstration that the trade-off between pulse duration and optical power approximates Bloch’s law, provides a rule of thumb, in both basic and preclinical studies^33^, for adjusting one of these parameters when the other has been changed. Moreover, our findings show that pulse duration can substitute effectively for optical power in controlling the size of the recruited population of opsin-expressing neurons. Computer control of pulse duration is readily implemented, whereas only a subset of optogenetic setups offer computer control of optical power.

High optical powers have been used to recruit neurons within a large region^34^. Although this is clearly useful over a limited range, the observed breakdown of reciprocity at the highest power tested suggests that this strategy may ultimately become futile in the case of ChR2. In order to recruit neurons within a large brain volume, a red-shifted opsin with high sensitivity^35^ offers a more promising approach.

Our results and methodological approach illustrate how the widespread use of a fixed set of stimulation parameters in many optogenetic behavioral studies unnecessarily adds across-subject variance and limits the inferences that can be drawn about underlying physical and physiological processes. Note that the pulse durations required to support ***TA_50_*** vary over more than a ten-fold range across stimulation sites at 20 mW and 31.6 mW (Appendix B). Consequently, the 0.5 ms pulse duration drove time allocation to near-maximal values in Rat OP16 at 20 mW but yielded minimal time allocation in Rat OP13 (Figure 3). This variance obscures the fact that the functional relationship between time allocation and the pulse duration in these two rats (and the others) is very similar; the same sigmoidal function fits the data very well. Thus, the congruence of the data from these two rats becomes evident when behavioral equivalence is used as the criterion for setting the values of stimulation parameters individually for each stimulation site but not when a fixed, informal standard (e.g., a pulse duration of 5 ms) is imposed. By sweeping the pulse duration over a large range for every rat, the sigmoidal form of the psychometric function is revealed. The fitted functions can then be normalized as was done here to reveal how the pulse duration and the optical power interact. Such normalization compensates for inevitable across-subject variation in the location of the optical-fiber tip and the expression of ChR2. Thus, we advocate customization of stimulation parameters for each subject to achieve behavioral or functional equivalence.

The findings of this study illustrate that constant-behavioral-output trade-off functions are typically much more informative of underlying physical and physiological processes than simple input-output relationships between the value of a stimulation parameter and the corresponding vigor of the stimulation-induced behavior^22^. The correspondence between the data reported here and Bloch’s law reflects such a trade-off. The underlying logic “sees through” multiple neural stages that translate neural activation into behavior so as to reveal quantitative properties of the processes underlying optogenetic excitation.

## Appendices

Appendix A: supplementary figures A1 and A2.

Appendix B: fitted parameter and d50 values.

## Acknowledgements

Stephen Cabilio developed and maintained the experimental-control and data acquisition software. The experimental-control and data acquisition hardware was designed, built, and maintained by David Munro. Jean Colombel developed the curve-fitting code. Karl Deisseroth and Ilana Witten kindly provided TH::Cre sires to establish our breeding colony.

## Author Contributions

**Vasilios Pallikaras**: Conceptualization, Formal analysis, Investigation, Methodology, Software, Visualization, Writing - original draft, Writing - review & editing. **Francis Carter:** Formal analysis, Investigation, Software, Visualization, Writing - review & editing. **David Natanael Velázquez Martínez:** Investigation, Methodology, Software, Writing - review & editing. **Andreas Arvanitogiannis:** Conceptualization, Resources, Supervision, Writing - review & editing. **Peter Shizgal:** Conceptualization, Formal analysis, Funding acquisition, Methodology, Project administration, Resources, Software, Supervision, Visualization; Writing - review & editing.

