## Appendices for "The trade-off between pulse duration and power in optical excitation of midbrain dopamine neurons approximates Bloch’s law"

### Appendix A

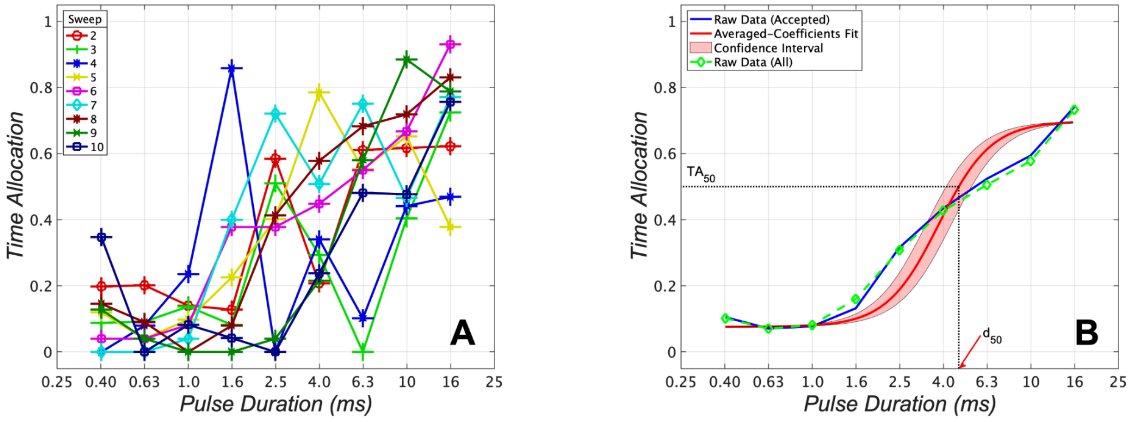


**Figure A1.** Pulse-duration sweep data for subject ELOP18 (optical power: 20 mW). **A:** Single-session data depicting individual time-allocation-versus-pulse-duration curves. **B:** Conventional averaging of time allocation over five test sessions (45 sweeps; green line) and curve produced by averaging the parameters of fitted sigmoidal functions (red line) with 95% confidence interval surrounding the location parameter (pink).

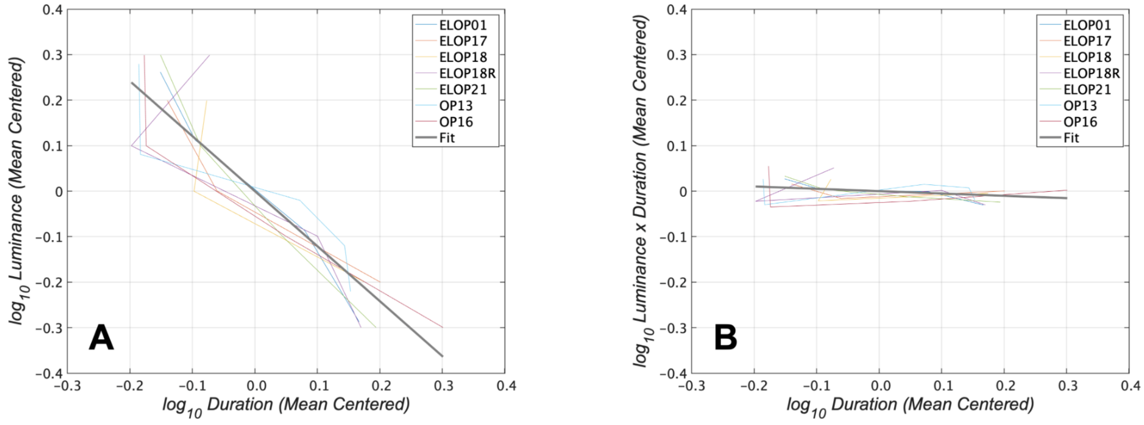


**Figure A2.** Mean-centered trade-off functions with data from all optical powers included. **A:** Mean-centered optical-power-vs-pulse-duration curves for individual stimulation sites (colored lines) and regression line (thicker black line; slope; -1.208 ± 0.236. **B:** Mean-centered energy (optical power × pulse duration) versus pulse-duration curves for individual stimulation sites (colored lines) and regression line fitted to the entire dataset (thicker black line; slope -0.0513 ± 0.064).

Appendix B: TAmax

| rat/power | 10mW | 12.6mW | 15.85mW | 20mW | 31.6mW | 44.6mW | 50mW |
|---|---|---|---|---|---|---|---|
| ELOP01 | | 0.837 | | 0.908 | 0.862 | 0.853 | |
| ELOP17 | | 0.852 | | 0.927 | 0.916 | | |
| ELOP18 | | | | 0.698 | 0.838 | | 0.807 |
| ELOP18R | | 0.832 | | 0.867 | 0.902 | | 0.894 |
| ELOP21 | | 0.752 | | 0.754 | 0.740 | | 0.667 |
| OP13 | 0.901 | 0.972 | 0.891 | 0.940 | 0.932 | | |
| OP16 | | 0.682 | | 0.634 | 0.705 | | 0.629 |

Appendix B: TAmin

| rat/power | 10mW | 12.6mW | 15.85mW | 20mW | 31.6mW | 44.6mW | 50mW |
|---|---|---|---|---|---|---|---|
| ELOP01 | | 0.025 | | 0.033 | 0.140 | 0.085 | |
| ELOP17 | | 0.091 | | 0.098 | 0.155 | | |
| ELOP18 | | | | 0.076 | 0.101 | | 0.124 |
| ELOP18R | | 0.070 | | 0.084 | 0.157 | | 0.059 |
| ELOP21 | | 0.054 | | 0.058 | 0.085 | | 0.092 |
| OP13 | 0.066 | 0.044 | 0.053 | 0.102 | 0.054 | | |
| OP16 | | 0.027 | | 0.078 | 0.104 | | 0.067 |

Appendix B: LocPar

| rat/power | 10mW | 12.6mW | 15.85mW | 20mW | 31.6mW | 44.6mW | 50mW |
|---|---|---|---|---|---|---|---|
| ELOP01 | | 0.491 | | 0.425 | 0.265 | 0.191 | |
| ELOP17 | | 0.514 | | 0.277 | 0.214 | | |
| ELOP18 | | | | 0.579 | 0.382 | | 0.396 |
| ELOP18R | | 0.607 | | 0.547 | 0.282 | | 0.377 |
| ELOP21 | | 0.629 | | 0.473 | 0.343 | | 0.216 |
| OP13 | 0.446 | 0.447 | 0.362 | 0.126 | 0.112 | | |
| OP16 | | -0.236 | | -0.504 | -0.692 | | -0.736 |

Appendix B: Slp

| rat/power | 10mW | 12.6mW | 15.85mW | 20mW | 31.6mW | 44.6mW | 50mW |
|---|---|---|---|---|---|---|---|
| ELOP01 | | 9.560 | | 9.030 | 8.672 | 9.376 | |
| ELOP17 | | 8.080 | | 9.240 | 8.387 | | |
| ELOP18 | | | | 8.354 | 10.244 | | 8.839 |
| ELOP18R | | 10.632 | | 8.650 | 8.633 | | 8.755 |
| ELOP21 | | 8.199 | | 7.300 | 7.265 | | 6.456 |
| OP13 | 10.441 | 11.410 | 12.882 | 11.020 | 11.207 | | |
| OP16 | | 14.470 | | 14.052 | 14.332 | | 13.889 |

Appendix B: d50

| rat/power | 10 mW | 12.6 mW | 15.85 mW | 20 mW | 31.6 mW | 44.6 mW | 50 mW |
|---|---|---|---|---|---|---|---|
| ELOP01 | | 0.527 | | 0.440 | 0.264 | 0.208 | |
| ELOP17 | | 0.532 | | 0.271 | 0.192 | | |
| ELOP18 | | | | 0.670 | 0.398 | | 0.419 |
| ELOP18R | | 0.632 | | 0.562 | 0.264 | | 0.389 |
| ELOP21 | | 0.699 | | 0.549 | 0.418 | | 0.354 |
| OP13 | 0.454 | 0.445 | 0.372 | 0.119 | 0.115 | | |
| OP16 | | -0.171 | | -0.422 | -0.646 | | -0.649 |
